# Classifying tumors from broad category to hierarchical molecular subtype: an interpretable hierarchical framework for precision oncology

**DOI:** 10.64898/2026.10.30.761789

**Authors:** Ǫingsong Zhang, Fei Liu, Xin Lai

## Abstract

Precise tumor subtype classification is important for guiding personalized treatment, characterizing tumor heterogeneity, and investigating molecular mechanisms. However, most existing multi-label classification models use flat architectures and do not fully exploit the inherent hierarchical relationships and biological constraints among tumor types, which limits their performance and interpretability in complex, fine-grained pan-cancer classification. Here, we present the cancer Hallmark-based Hierarchical Recursive Model (HHRM) for pan-cancer hierarchical multi-label classification. HHRM integrates hierarchical label structures with recursive neural networks, enabling simultaneous modeling of multi-level dependencies from broad cancer categories to specific subtypes. We trained the model on a comprehensive pan-cancer cohort of 11,476 samples spanning 26 primary cancer types and 405 subtypes across up to eight hierarchical levels. HHRM achieved an accuracy of 0.7924 for predicting hierarchical tumor subtypes and was further evaluated on a validation dataset of 887 samples. Compared with other hierarchical modeling frameworks, HHRM achieved higher accuracy in local, global, and mixed output modes. Ablation experiments showed that replacing the hallmark-based graph convolutional network reduced model performance. On an independent dataset of 1,608 breast cancer samples, HHRM also achieved an accuracy greater than 0.86. Finally, we combined SHAP with a network-based method to rank and analyze predictive genes at different levels of the tumor hierarchy, which identified both known and potentially novel biomarkers for cancer diagnosis. In conclusion, to our knowledge, HHRM is the first hierarchical model for multi-level tumor subtype classification. Its competitive performance, combined with post hoc biological interpretation, suggests potential value for precision cancer diagnosis.

**Availability and implementation:** The data, source code, and documentation of the study are available at the GitHub repository https://github.com/laixn/HHRM.

## Introduction

The classification of malignant tumors has evolved from traditional histological methods to precision systems based on multidimensional molecular features, including genomics, transcriptomics, and epigenomics. Due to the widespread adoption of high-throughput sequencing and international initiatives such as The Cancer Genome Atlas (TCGA), researchers have identified hundreds of tumor subtypes with unique molecular properties [1], [2]. These subtypes differ in clinical prognosis, treatment response, pathogenic mechanisms, and therapeutic targets [3], [4]. In clinical practice, accurate identification of tumor subtypes directly informs treatment decisions. For example, molecular subtypes of breast cancer (Luminal A/B, HER2-positive, and triple-negative) influence the choice between endocrine therapy, targeted therapy, and chemotherapy [5]. Additionally, the tumor subtype classification system can help diagnose cancers with unknown primary origins, which account for 2%-5% of all malignancies and are the fourth leading cause of cancer-related death [6], [7]. Despite their clinical importance, current cancer subtype classification systems have two structural limitations. First, subtype systems for different cancer types have been developed independently without a unified framework. Second, the biological hierarchy connecting broad cancer categories to increasingly specific subtypes has not been systematically modeled. This constrains the biological consistency and clinical applicability of existing classification schemes.

These limitations are exacerbated by the challenges inherent in computational subtype classification, which is essentially a multi-class pattern recognition problem in a high-dimensional feature space. First, tumor multi-omics data typically involves tens to hundreds of thousands of features, such as gene expression, mutations, and copy number variation, but only thousands to tens of thousands of samples [8]. This produces a “high-dimensional, small-sample” problem that is further complicated by technical heterogeneity across cancer types, sequencing platforms, and batches [9]. Second, tumor subtype systems are hierarchical. For example, the classification of lung adenocarcinoma progresses from the broad category of lung cancer to a specific tissue or cell type, such as adenocarcinoma, squamous cell carcinoma, large cell carcinoma, or non-small cell carcinoma, and finally to a molecular subtype, such as EGFR-mutant or ALK-fusion [10]. This hierarchical relationship shows important biological constraints, where biological entities shared at higher levels may undergo specific alterations at lower levels. Third, subtype classification systems are dynamic rather than fixed [4]. New molecular subtypes continue to be reported, and many rare subtypes have limited sample sizes, challenging the generalization of any classifier trained on them.

Existing computational approaches only partially address these challenges. Flat classification methods treat subtype prediction as a standard binary or multi-label problem, failing to model hierarchical relationships [11], [12], [13], [14], [15]. Although these methods perform well when the label space is small and shallow, usually limited to primary types [16], [17] and secondary subtypes [18]. Pan-cancer subtyping can span labels across multiple hierarchical levels in much deeper hierarchy [19]. Consequently, the performance of flat classifiers tends to decline as the number of labels increases, partly because they ignore the molecular continuity linking related subtypes. Although hierarchical classification methods explicitly model subtype structure, existing implementations have significant limitations. One approach stratifies samples by cancer type and classifies each type independently rather than across a unified, pan-cancer cohort. Its hierarchy lacks a clear clinical basis [19], [20]. Another approach classifies non-TCGA samples according to TCGA-derived molecular subtypes. However, its hierarchy is limited to two levels, making it difficult to capture cross-cancer and cross-level relationships that only emerge in deeper hierarchies [18].

Convolutional neural networks (CNNs) and graph neural networks (GNNs) have been used for tumor classification. CNNs are primarily used for medical imaging or image-transformed omics data [21], while GNNs are used to incorporate tissue structure or gene interaction networks via pathological graphs [22], [23] or gene networks [24], improving performance and interpretability. However, these methods usually target a single cancer type or data modality, lacking a unified framework for multi-level, pan-cancer classification. Hierarchical classification is well-established outside of oncology, for example, in protein function prediction [25], [26], gene ontology annotation [25], [27], cancer cell line annotation [28], and microbiome clustering [29]. These applications build on general hierarchical classification methodology [30]. However, to date, this approach has not been applied to pan-cancer tumor subtype classification with multiple levels.

Taken together, no existing computational framework spans a unified pan-cancer cohort, models the full, multi-level, hierarchical structure of tumor subtypes, and grounds its predictions in interpretable biological mechanisms, even though hierarchical modeling is a mature approach in adjacent biomedical domains. Therefore, a tool for accurate hierarchical tumor subtype classification can improve current cancer diagnostics and help resolve malignancies of unknown origin, both are persistent challenges central to precision cancer medicine.

Toward this goal, we develop the cancer Hallmark-based Hierarchical Recursive Model (HHRM), a computational framework for pan-cancer, hierarchical, multi-label classification (Figure 1). The HHRM performs hierarchical transfer and refinement of feature representations through a recursive mechanism. Higher levels learn shared feature representations of broad cancer categories, and lower levels build on this foundation to learn subtype-specific features. This forms a coarse-to-fine decision process. The model’s design is based on cancer hallmark gene networks and addresses the biological constraint that information shared at higher hierarchical levels is refined rather than discarded at lower levels. We train and evaluate HHRM on a recent pan-cancer dataset containing 26 cancer types, 405 hierarchical molecular subtypes across up to eight levels, and over 10,000 samples. HHRM achieves an average accuracy of 0.7924 while preserving hierarchical constraints, outperforming existing hierarchical models. To assess generalization, we apply the model to an independent breast cancer dataset containing 1,608 samples, four subtypes, and two levels, achieving an accuracy of over 0.90. Finally, we use SHAP and PathwaySpace to provide a quantitative, pathway-level biological interpretation of the model’s predictions and offer a framework for understanding its hierarchical decision-making mechanism.

**Figure 1.**
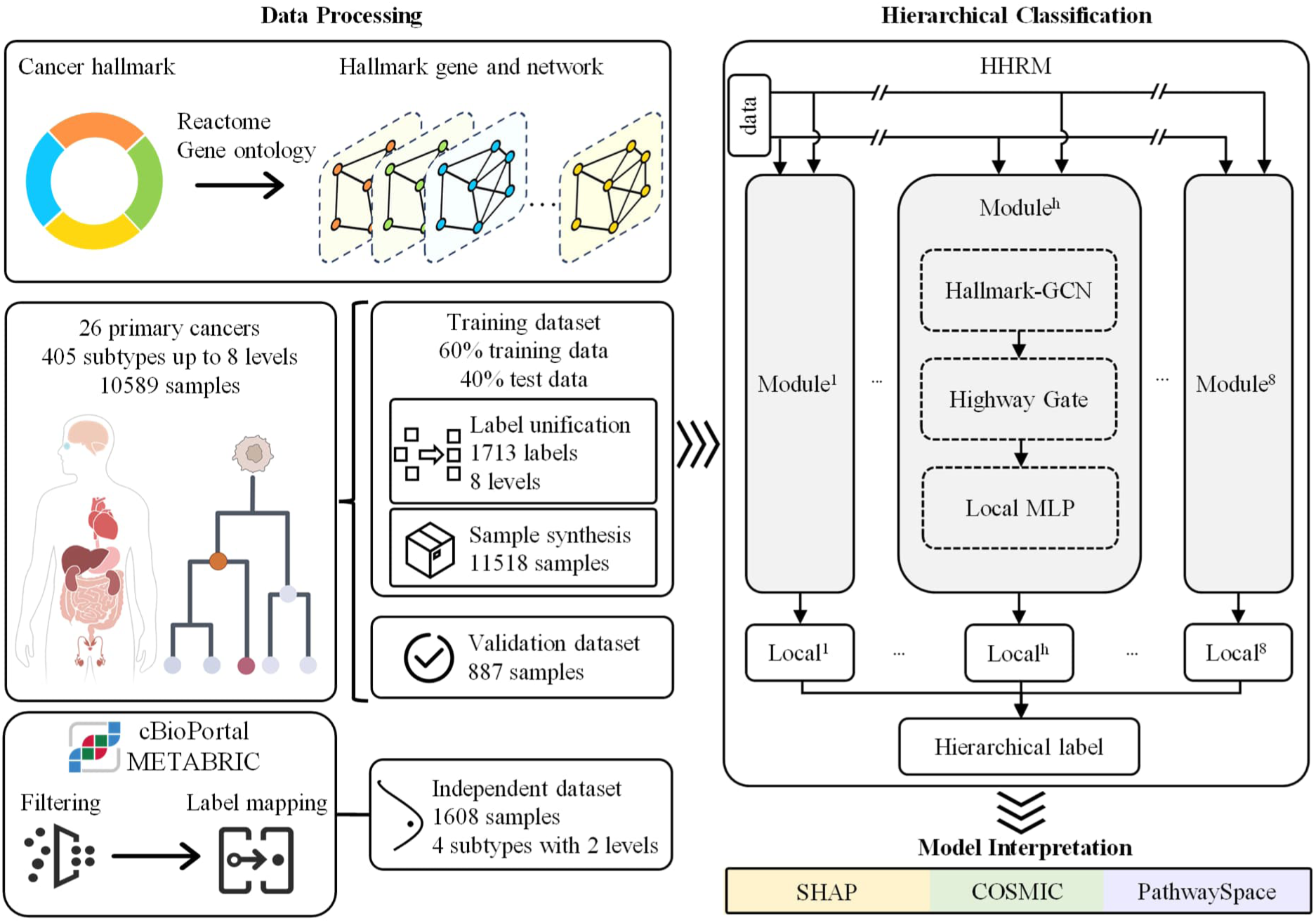
Overview of the study workflow. First, the training and independent datasets are processed, and the cancer hallmark networks are reconstructed. Second, the cancer hallmark-based hierarchical recursive model (HHRM) is developed to classify hierarchical tumor subtype labels. Finally, SHAP, PathwaySpace and the COSMIC database are employed to quantify the contribution of genes to the model’s predictions at both gene and hallmark levels and examine their association with known cancer driver genes.

## Methods

### 2.1 Data Collection and Processing

We downloaded gene expression data from the Treehouse data (v11) (https://xena.treehouse.gi.ucsc.edu:443), a comprehensive cohort that collects and annotates more than 11,000 patient samples from TCGA and TARGET. We obtained samples’ original hierarchical labels from [19] and processed them as described in our previous work [31], which classified tumors at different stages based on morphological and genomic information (Figure 2). Sample labels consist of an identifier and an abbreviated tumor origin site (e.g., T000 CNS for central nervous system cancer), followed by subtype classification based on histological features and genetic mutations (e.g., T028 CNS IDHwt and T029 CNS IDHmut for CNS cancer with IDH1-wildtype and IDH1-mutation, respectively). Each label may span up to eight hierarchical levels. Level 1 denotes the primary tumor type, and each subsequent level denotes a subtype of the level above it, with a given level potentially containing multiple labels. For example, the primary-level leukemia label (T005 LEUK) has two level-2 subtypes, acute lymphoblastic leukemia (T119 ALL) and acute myeloid leukemia (T120 AML), which, based on morphological phenotype and genetic mutation, extend to up to 8 and 4 levels, respectively. For the 26 tumor types in this cohort, we obtained subtype markers spanning levels 1 to 8 (Supplementary Excel Sheet S1).

**Figure 2.**
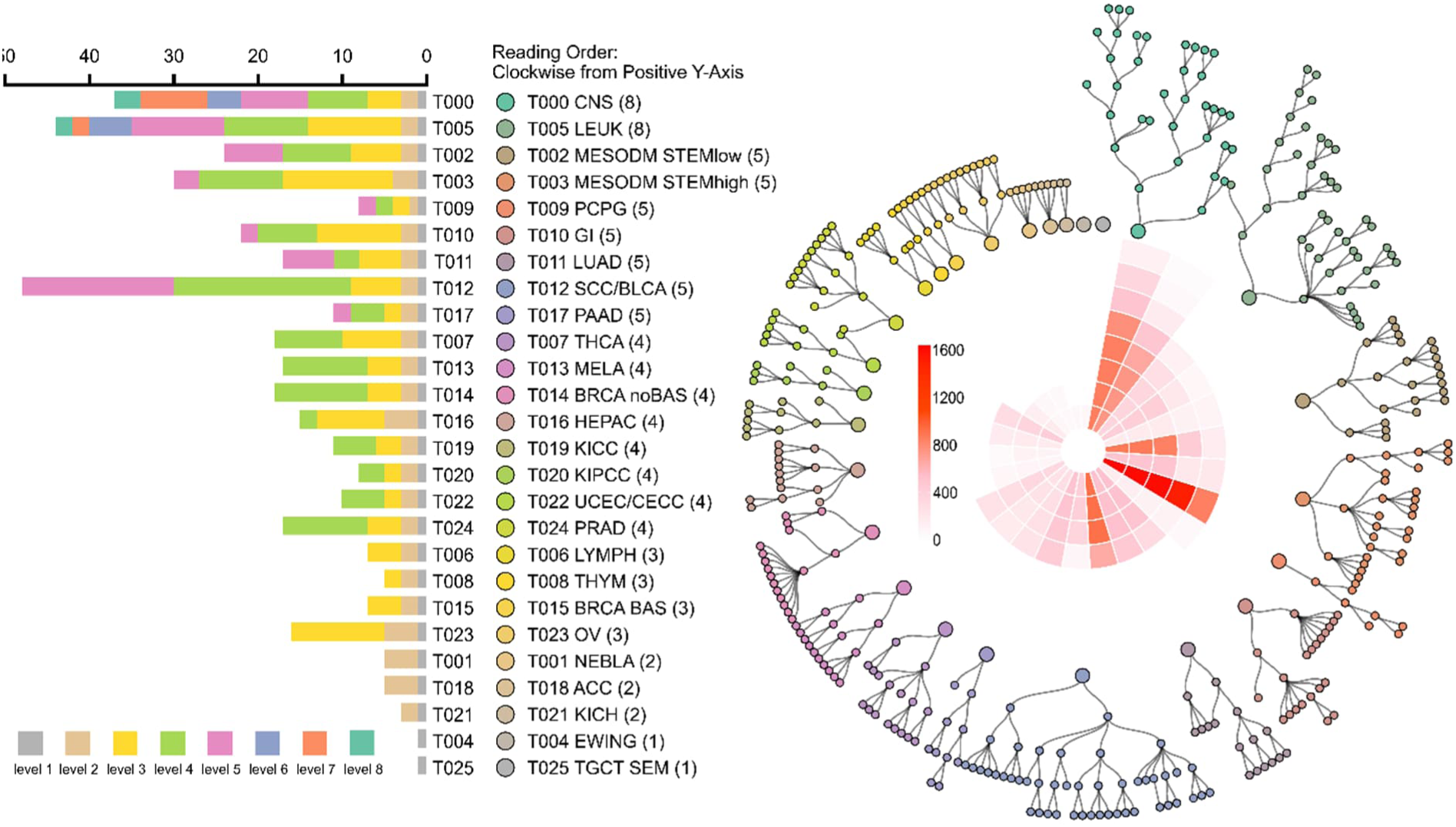
Overview of the Treehouse data. The bar plot shows the number of labels at different levels contained within 26 cancer types. The tree plot shows the hierarchical structure of labels in each cancer type, as well as the corresponding number of samples at each level. The darker the red, the more samples correspond to that level of the hierarchy. The circles in both plots use the same color code to represent different types of cancer.

After curation, we obtained a pan-cancer cohort of 11,476 samples spanning 26 primary cancers and 405 tumor subtypes, with hierarchies of up to 8 levels. We then divided these samples into TRUE and FALSE groups following the rule in our previous work [31]. For each sample, we searched for cancer types that aligned with the clinical diagnosis across the sample’s label levels. Samples with matching types across all label levels were categorized as TRUE, and all others as FALSE. This resulted in 10,589 TRUE samples as a training dataset that is further expanded by adding synthetic samples and 887 FALSE samples as a validation dataset.

To standardize the hierarchical depth of the TRUE samples, we unified their label levels (Figure S1). For each TRUE sample with fewer than 8 label levels, we expanded its labels to 8 levels using synthetic labels with no biomedical meaning, by replicating the label of its deepest real level. For example, a sample with a 4-level label (T005 LEUK - T120 AML - T143 AMKL - T156 AMKL HOX) was expanded from level 5 onward by replicating its level-4 label (T156 AMKL HOX_syn_5 - T156 AMKL HOX_syn_6 - T156 AMKL HOX_syn_7 - T156 AMKL HOX_syn_8). This procedure increased the number of unique labels at each level from [26, 56, 114, 126, 59, 9, 10, 5] to [26, 64, 155, 253, 295, 301, 307, 312], for a total of 1,713 unique labels across the eight levels among the 10,589 TRUE samples, each sample retaining one label per level (Supplementary Excel Sheet S2).

The distribution of samples across cancer types is uneven. Some common cancers, such as lung and skin cancers, have large sample sizes, while some rare cancers, such as bone cancer and sarcoma, have limited samples. To address this imbalance, we generated synthetic data for labels with fewer than 20 samples, so that every label had at least 20 samples. For labels with more than one existing sample, we used linear interpolation to assign values to all features based on the maximum and minimum feature values of the existing samples. For labels with only a single corresponding sample, we generated additional samples by adding random noise using the equation, *synthetic_value* = 0.1 × *sign* × *feature_value* × *noise*, where noise followed a normal distribution (*μ* = 0, *σ* = 1) and sign was randomly assigned as −1 or 1. Synthesizing data for the 109 labels with fewer than 20 samples expanded the model training dataset to 11,518 samples.

### 2.2 Cancer hallmark genes and networks

As described in our previous work [31], we created ten cancer hallmark gene sets using relevant GO terms, resulting in 6,448 genes. We then downloaded 75,869 molecular interactions from the Reactome database [32] and used these interactions to reconstruct ten hallmark-related gene regulatory networks. These hallmark networks were used for model feature embedding via a graph convolutional network (Method Section 2.3).

### 2.3 Cancer hallmark-based hierarchical recursive model

Inspired by a published machine learning framework in computer science [33], we developed a hierarchical recursive model to perform end-to-end multi-label classification for hierarchical tumor subtypes (Figure 3). The model’s input are 6,448 genes’ expression levels (i.e., log₂(TPM+1)) and ten hallmark networks. The model consists of eight recursive modules, with each module outputting a local output for one hierarchical level. Each module is composed of three units that are a cancer hallmark-based graph convolution network (Hallmark-GCN) for feature embedding, a highway gate for information filtering, and an output unit for predicting tumor subtypes at the current level.

**Figure 3.**
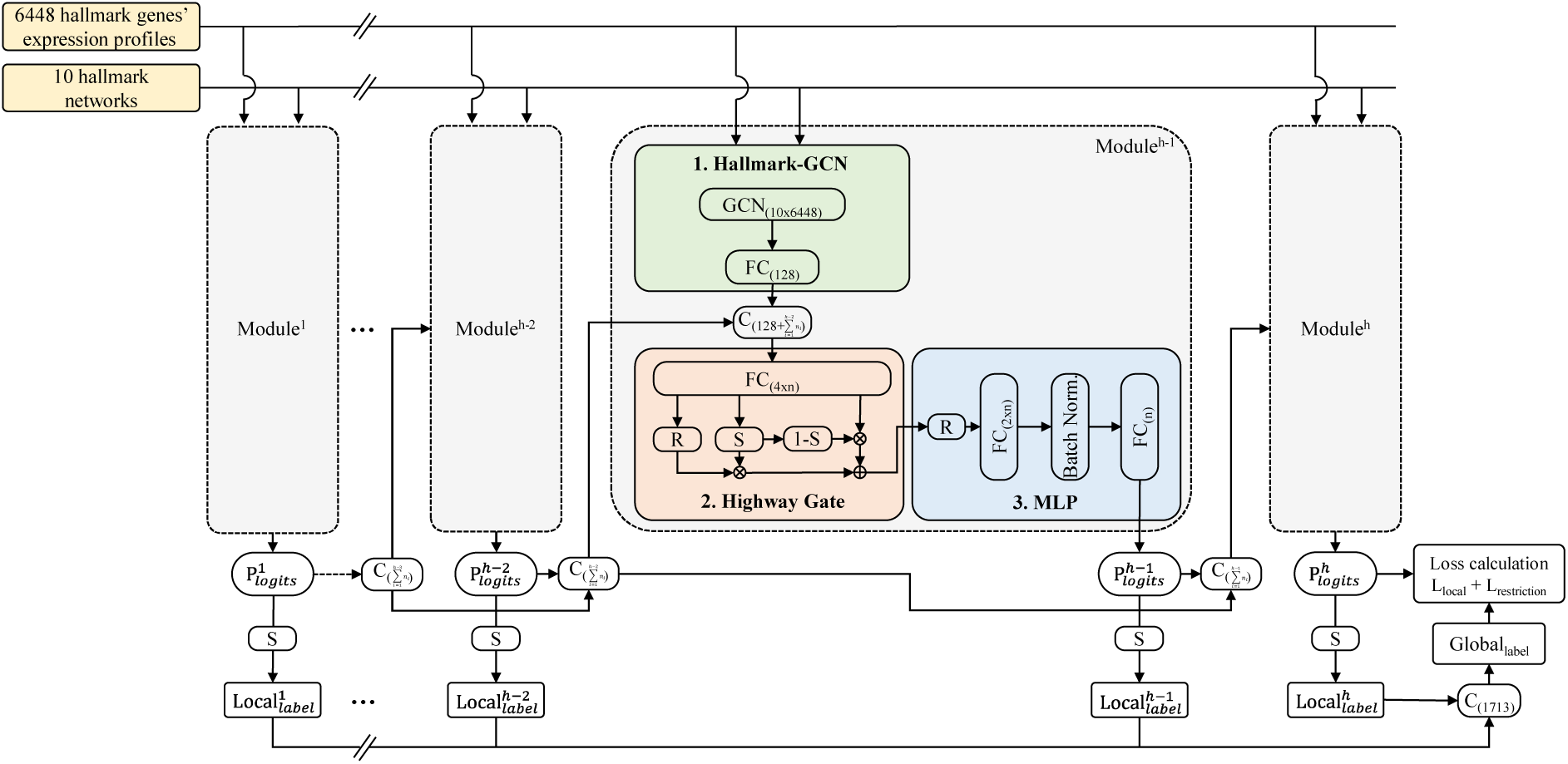
The architecture of the model. The input data is 6,448 genes’ expression levels and ten hallmark networks. The model contains eight recursive modules, and each module has three units including a Hallmark-GCN for feature embedding, a highway gate for filtering information, and an MLP for predicting local labels. Specifically, Hallmark-GCN performs graph convolution to model gene-gene message passing over the undirected hallmark network. The highway gate captures categorical dependencies across hierarchical levels by filtering current-level data with predictions from the previous level. The MLP comprises two fully connected layers with batch normalization. To make hierarchical label prediction, the predicted local labels from the current and previous HHRM modules together with the input data are sent into the next module. Finally, the local labels are concatenated to form the global label. *R*, *S*, and *C* indicate ReLU function, sigmoid function, and concatenation, respectively. The numbers and equations in parentheses show the dimensions of the data. *n* indicates the number of labels at the current level, with a maximum of eight levels (*h* = 8).

The Hallmark-GCN embeds information from gene expression profiles and the hallmark networks at the corresponding level, *Data_l_^h^* using the following equation, in which *l* and *h* indicate the *l_th_* round of information aggregation using hallmark network and *h*_th_ layer in the hierarchical labels, respectively.

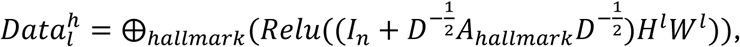

Where *H^l^* is the input matrix for the *l*_th_ round of information aggregation using GCN, and *W^l^*is the learnable weight matrix for the *l*_th_ round and has the same size as *H^l^*. We used ten hallmark networks (*l* = 1,…,10), and the GCN aggregates each gene’s own expression with that of its direct neighbors in each network.

This is denoted by the term 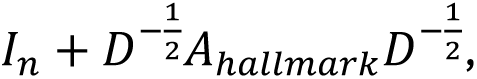 which contains the hallmark network’ adjacency matrix *A_hallmark_*, the identity matrix *I_n_*, and the degree matrix *D*, corresponding to each gene’s node degree in the undirected hallmark network. *ReLU* is a non-linear activation function that sets all negative values to zero. Finally, ⨁_ℎ*allmark*_ denotes concatenation of the output computed using the same procedure for all cancer hallmarks, resulting in a matrix comprising 6,448 genes, 11,518 samples, and 10 cancer hallmarks. The matrix passes through a fully connected layer of 128 neurons to reduce the dimensions of the features.

The matrix *Data_l_^h^* is passed to the highway gate unit for information filtering [33], [34]. This unit models the categorical dependencies between hierarchical tumor subtypes at different levels by filtering the current level’s data using prediction information from the preceding level. Specifically, the input matrix for the highway gate unit at level *h* is constructed as

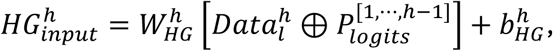

where *W_HG_^h^* is a learnable weight matrix, *b^h^_HG_* is a bias vector, and ⊕ denotes concatenation of *Data^h^_l_* (the Hallmark-GCN output at the current level *h*) with *P_logits_*^[1,…,*h*-1]^ (the label predictions from levels 1 through h−1). The unit’s input has a length of 128 plus the number of cancer subtypes from all preceding levels. This input is connected to a fully connected layer with 4×*n* neurons, where *n* is the number of cancer subtypes at the current level, so that the number of neurons scales dynamically with the number of labels at each level. The data is then further filtered using the equation

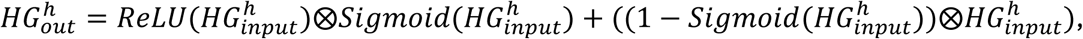

where ReLU and Sigmoid are activation functions, and ⊗ denotes matrix multiplication.

The highway gate’s output is then passed to an MLP unit for predicting tumor subtypes at the current level. This unit contains two fully connected layers and a batch normalization layer; as with the first fully connected layer in the highway gate unit, their sizes are 2×*n* and *n*, respectively. The batch normalization layer stabilizes and accelerates training by normalizing hidden-layer activations to zero mean and unit variance within each mini-batch.

The model’s output consists of eight local predictions (*Local_label_^h^*; *h* = 1…8), obtained by applying a Sigmoid activation to the model’s raw output (*P_logits_^h^*) at each level. These local predictions are concatenated into a global prediction (*Global_label_*) representing a sample’s hierarchical label from level 1 to level 8.

### 2.4 Loss Function

We designed a loss function, termed weighted focal cross entropy loss. This function considers three aspects, including sample imbalance across cancer types, local predictions of cancer subtypes at each hierarchical level, and the global prediction of the full hierarchical label.

To address label imbalance, we first calculated a weight for each label at each level, proportional to the number of corresponding samples, so that labels with fewer samples received higher weights and labels with more samples received lower weights. The weight of label *i* at level *h* was computed as:

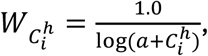

where *C_i_^h^* is the label’s count at the level *h* and *a* is the coefficient, set to 1.02. We then normalized labels’ weight at each level

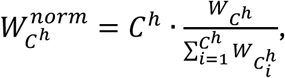

where *W C^h^* is a label’s weight at level *h* and *C^h^* is the total number of labels at level *h.* This normalization ensures that the sum of normalized label weights at level h equals the number of labels at that level.

We next used the following equation to compute the weighted, label-smoothed cross-entropy loss (*Loss_CEsmoot_*_ℎ_)

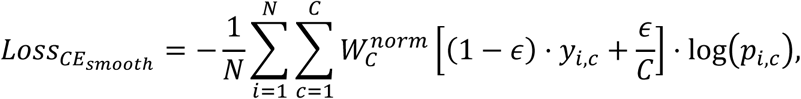

where *N* is the number of samples and *C* is the total number of labels. *y* is a vector of 1,713 values (the total number of labels), with a value of 1 for the sample’s true label and 0 otherwise; *p* represents the samples’ raw predictions (i.e., logits) prior to sigmoid normalization. The smoothing coefficient ε was set to 0.3.

We then calculated the focal cross-entropy loss for each level

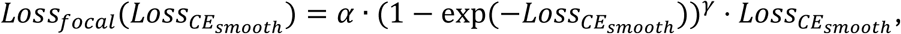

where *α* and *γ* are coefficients that are set to 1.0 and 2.0, respectively.

Finally, the total loss (*L_all_*) was defined as the sum of a local loss (*L_local_*) and a restriction loss (*L_restriction_*).

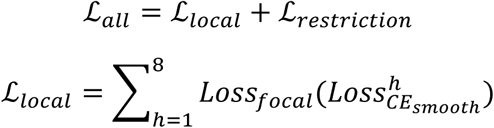

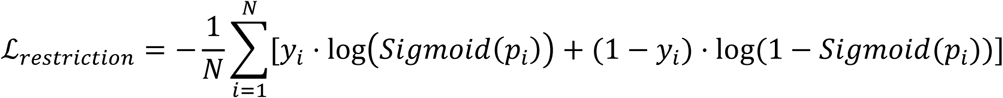

*L_local_* is the sum of the focal cross-entropy losses across all eight levels. *L_restriction_* is the multi-label cross-entropy loss computed on the concatenated local predictions across all levels, this term adds hierarchical restriction on local labels, following the label-unification rule described in Method Section 2.1. In the equation, *y_i_* and *p_i_* are the true label vector and logits for sample *i*, respectively.

### 2.5 Model training and validation

For model training, we split the 11,518 samples into a training set (60%) and a test set (40%) using multiple random seeds (Figure S2). To maintain the proportion of labels from different cancer types at each level, as in the original data, we performed hierarchical sampling at each level. We selected the best-performing seed and trained the model from five different initializations. We used mean accuracy ± standard deviation to report the model’s performance on the test data. We evaluated the best model with the highest accuracy on the 887 validation samples that were excluded from model training. Model performance on the validation data was assessed using four metrics: accuracy, macro-precision, macro-recall, and macro-F1. During training, we used a mini-batch gradient descent approach (batch size = 200) to improve computational efficiency.

We additionally evaluated HHRM and the benchmark models on an independent dataset METABRIC. The microarray dataset contained three breast cancer cohorts [35], [36], [37] and was downloaded from cBioPortal (https://www.cbioportal.org). After processing and filtering, this dataset contained 1,608 valid samples with four breast cancer subtypes (i.e., Basal, Lum A, Lum B, and HER2). We transformed these four labels into hierarchical labels with two levels that are related to TCGA subtypes. This allowed us to retrain models for classifying these tumor subtypes (Supplementary Excel Sheet S3). Specifically, Basal was transformed into a single label that included two TCGA Basal subtypes. The Lum A, Lum B, and HER2 subtypes were transformed into two labels. The dataset contains 6,084 expressed hallmark genes, while the expression value of other unexpressed hallmark genes is set to zero. The expression data were transformed using the z-score method. We split the METABRIC data into 60% for training, 20% for testing, and 20% for validation, and reported model performance on this split.

### 2.6 Model benchmarking

We compared HHRM with several state-of-the-art models used for hierarchical multi-label classification (Table S1). All benchmark models were trained and evaluated under identical conditions, including the same data split and default hyperparameter settings (Supplementary Excel Sheet S4). Because the benchmark models have different output modes for predicting hierarchical labels, we evaluated each model under three output modes to ensure fair comparisons (Figure S3).

(I) Local mode combines all local labels according to the label-unification rules to form a sample’s hierarchical label. Each of the eight local outputs contains the number of labels at its corresponding level (e.g., 26 labels at level 1, 253 labels at level 4), and the local label with the maximum probability is selected as that level’s prediction. This is the original output mode of HHRM. The term *L_restriction_* in HHRM’s loss function *L_all_* = *L_local_* + *L_restriction_* ensures that the predictive combined local labels match with samples’ true hierarchical labels.
(II) Global mode directly outputs samples’ hierarchical label. Since HHRM lacks a global layer, we added one to make its predictions comparable to those of benchmarking models containing a global layer for hierarchical predictions. Accordingly, we added a new loss term, *L_global_*, to HHRM’s original loss equation: *L_all_* = *L_local_* + *L_restriction_* + *L_global_*. The new term has the same mathematical definition as *L_restriction_*. The new term allowed HHRM to predict 1,713 labels for each sample simultaneously. Predicted labels with probability exceeding a threshold of 0.5 (adapted from HARNN [33]) are used to reconstruct a sample’s hierarchical label.
(III) Mixed mode combines the predictions from the local and global modes by computing a sample’s prediction as *β* × *local* + (1−*β*) × *global*, with *β* = 0.5, giving equal weight to each component. More specifically, the equation uses a combination of predictions from the local layers and the hierarchical label predicted by the global level to determine the final output. As in global mode, predicted labels with a probability greater than 0.5 are used to reconstruct a sample’s hierarchical label.

For all three output modes, a predicted hierarchical label was regarded as correct only if it matched the sample’s predefined label at all eight levels; otherwise, it was regarded as incorrect.

### 2.7 Feature ranking and interpretation using a network-based approach

We used SHapley Additive exPlanations (SHAP) to characterize the importance of genes for our model [38]. We randomly selected 1,000 samples from the complete dataset to compute the SHAP values of their features and ranked genes by their absolute SHAP values, with higher values indicating greater importance to the classifier.

We used PathwaySpace to map the cancer hallmark genes onto the gene regulatory network derived from Pathway Commons (v12), comprising 12,990 genes and 359,540 interactions from 22 databases [39]. PathwaySpace converts this network into a two-dimensional landscape to quantify signal propagation and enrichment patterns across the network [18]. In this framework, each gene’s SHAP value serves as its initial signal intensity, designating it as a signal source; signals propagate outward from each source along network edges, attenuating according to the Weibull decay function (PathwaySpace’s default attenuation function). At each point in the 2D landscape, the attenuated signals from neighboring genes are aggregated into a single scalar using weighted mean aggregation, appropriate for continuous signal values such as SHAP values, yielding a density value at that point. Higher density values reflect greater local enrichment of high-SHAP genes and stronger signal convergence in the neighboring genes, whereas lower values indicate sparser gene signal enrichment.

Using the PathwaySpace results, we assessed the overlap between the top SHAP-ranked genes and COSMIC’s 554 tier 1 cancer driver genes [40] as well as their clusters. Cancer gene clusters were identified by performing heat diffusion and watershed segmentation on a 2D landscape, followed by gene set enrichment analysis to assign a thematic name to each cluster representing one of the 30 major cancer-associated pathways [18].

To annotate the association of 38 SHAP-ranked genes with cancer, we searched the literature using the academic AI search engine Consensus (https://consensus.app) with the prompt “identify the association of gene X with cancer”, where X was each gene symbol. We then retrieved each publication the tool returned and reviewed it against the original article to confirm that it supported the stated association. The verified evidence for each gene was summarized and classified into three categories. Genes were classified as having a strong association with cancer when direct experimental evidence showed that they were dysregulated in cancer, were associated with survival, and affected cancer phenotypes such as proliferation or apoptosis. Genes supported only by mutation data, without direct evidence of a functional role in cancer, were classified as having a very weak (1≥n≥2) or weak (n≥3) association, depending on the number of publications found. The full annotations are provided in Supplementary Excel Sheet S6.

## 3 Results

### 3.1 HHRM surpasses state-of-the-art methods on pan-cancer hierarchical classification

We developed HHRM to classify hierarchical tumor subtypes in a pan-cancer cohort. The cohort contains transcriptomic profiles of patients that are used for clustering patients, which results in tumor subtypes with a clinical basis [19]. A recent study that used machine learning models to classify non-TCGA cancer samples into TCGA molecular subtypes revealed the clinical need for hierarchical tumor subtype classification [18]. After preprocessing and cleaning, we obtained a dataset of 11,476 samples from 26 primary tumors. Each tumor sample has a label spanning up to eight hierarchical levels and was processed for classification tasks (Method Section 2.1).

To handle hierarchical tumor subtypes, HHRM has an architecture of recursive modules, each of which uses hallmark information to predict the corresponding hierarchical label of a sample (Method Section 2.2 and 2.3). To our knowledge, HHRM is the first machine learning model that uses cancer hallmark networks for feature embedding to predict hierarchical tumor subtypes. HHRM was inspired by HARNN a text classification model [33]. We modified its architecture in three respects for hierarchical multi-label classification. First, in addition to gene expression profiles, we used the local outputs from all preceding levels as input for each subsequent level. Second, we concatenated local outputs according to our label-unification rules to reconstruct the hierarchical label that is the model’s global output (Method Section 2.1). Third, we applied the weighted focal cross-entropy loss function to simultaneously mitigate the long-tail effect and consider hierarchical label constraints during model training (Method Section 2.4). Unlike our previous model, HallmarkGraph [31], which predicts labels at each level individually, HHRM directly predicts a sample’s hierarchical label up to its original label’s hierarchical level.

Because HHRM is specifically designed for hierarchical multi-label classification across pan-cancer subtypes, we chose several state-of-the-art models that are mainly applied in the domain of hierarchical multi-label classification as benchmarks (Table S1). HMLC_LMLP is a hierarchical model in which each level is trained and predicted independently using an MLP, without a global output (i.e., a connected hierarchical label) [26]. HMCN_F uses cascaded modeling (the output of each preceding level serves as input to the next level) with multiple local outputs and one global output [25]. HMCN_R is an extension of HMCN_F with a recurrent structure incorporating forget gates, input gates, and output gates [25]. HARNN applies hierarchical attention within the recurrent structure for cascaded modeling and produces multiple local outputs and one global output [33]. C-HMCNN enforces a mandatory hierarchical constraint matrix and has only a single global output [27]. We also used an MLP, which is a fully connected neural network with only a single global output and without any hierarchical constraint.

Since the benchmark models produce hierarchical labels differently, we compared HHRM with them under three output modes (local, global, and mixed; Method Section 2.6) to ensure a fair comparison. Each mode represents a distinct strategy for predicting a sample’s hierarchical label. HHRM outperformed all benchmark models across all four metrics in all three modes (Table 1). In the local mode, HHRM achieved an accuracy of 0.7924, which was 0.003-0.078 higher than those of the benchmark models. HMCN_R (0.7896) performed most similarly to HHRM, and HMCN_F (0.7142) performed worst. In the global mode, HHRM’s accuracy (0.7815) was 0.0156-0.0234 higher than the other benchmark models. HMCN_R (0.6257) had the second-best accuracy and HMCN_F (0.5471) the lowest. C-HMCNN and MLP, which have only a single global output, were evaluated in this mode only. In the mixed mode, HHRM’s accuracy (0.7520) was 0.011-0.0178 higher; HMCN_R (0.6420) was again second best, and HARNN (0.5742) was lowest. HHRM also ranked first in macro-precision, macro-recall, and macro-F1 in every mode. For example, its macro-F1 was 0.8460, 0.8372, and 0.8290 in the local, global, and mixed modes, compared with 0.8292 (HARNN), 0.7837 (MLP), and 0.7643 (HMCN_R) for the best-performing benchmark in each mode. These results indicate that HHRM has superior capability for extracting and filtering information for hierarchical tumor subtype classification.

**Table 1.** Benchmark results. The table shows the metrics for different models with three output modes. The metrics shown are the average performance of five individual runs and standard deviation. The best metric is bold, and the second-best metric is underlined.

| Output mode | Model | Accuracy | Macro-Precision | Macro-Recall | Macro-F1 |
|---|---|---|---|---|---|
| Local mode | HHRM (ours) | **0.7924±0.0040** | **0.8658±0.0014** | **0.8422±0.0027** | **0.8460±0.0022** |
|  | HARNN | 0.7443±0.0023 | 0.8505±0.0016 | 0.8241±0.0028 | 0.8292±0.0020 |
|  | HMCN_F | 0.7142±0.0107 | 0.8204±0.0056 | 0.7852±0.0028 | 0.7919±0.0024 |
|  | HMCN_R | 0.7896±0.0013 | 0.8525±0.0013 | 0.8206±0.0017 | 0.8268±0.0015 |
|  | HMC_LMLP | 0.7622±0.0020 | 0.8486±0.0014 | 0.8188±0.0008 | 0.8236±0.0008 |
| Global mode | HHRM (ours) | **0.7815±0.0032** | **0.9103±0.0399** | **0.8049±0.0018** | **0.8372±0.0012** |
|  | HARNN | 0.5583±0.0016 | 0.8838±0.0080 | 0.6883±0.0067 | 0.7509±0.0031 |
|  | HMCN_F | 0.5471±0.0287 | 0.8227±0.0091 | 0.6790±0.0183 | 0.7274±0.0115 |
|  | HMCN_R | 0.6257±0.0018 | 0.8614±0.0005 | 0.7232±0.0016 | 0.7708±0.0010 |
|  | HMC_LMLP | 0.5932±0.0014 | 0.8486±0.0012 | 0.7269±0.0019 | 0.7696±0.0012 |
|  | C-HMCNN | 0.6185±0.0017 | 0.8409±0.0022 | 0.7416±0.0027 | 0.7761±0.0010 |
|  | MLP | 0.5754±0.0023 | 0.7927±0.0040 | 0.7932±0.0048 | 0.7837±0.0025 |
| Mixed mode | HHRM (ours) | **0.7520±0.0031** | **0.9021±0.0015** | **0.7858±0.0029** | **0.8290±0.0021** |
|  | HARNN | 0.5742±0.0016 | 0.8962±0.0042 | 0.6963±0.0061 | 0.7631±0.0036 |
|  | HMCN_F | 0.5775±0.0265 | 0.8596±0.0103 | 0.6739±0.0201 | 0.7367±0.0125 |
|  | HMCN_R | 0.6420±0.0013 | 0.9009±0.0015 | 0.6974±0.0015 | 0.7643±0.0012 |
|  | HMC_LMLP | 0.6061±0.0013 | 0.8967±0.0010 | 0.6952±0.0008 | 0.7636±0.0006 |

### 3.2 Ablation of Hallmark-GCN reduces model performance

We conducted a series of ablation experiments to assess the contribution of individual components to the overall model performance (Table 2). We first tested two HHRM variants: one aggregating the local outputs of all preceding levels (inspired by DenseNet [41]) and the other aggregating only the local output of the immediately preceding level, with the gene expression profiles as input, for the subsequent module containing the Hallmark-GCN and highway gate. The two variants performed comparably. The variant using all preceding levels showed a similar accuracy (0.7981 vs. 0.7921), whereas its macro-precision (0.8585 vs. 0.8677), macro-recall (0.8284 vs. 0.8396), and macro-F1 (0.8342 vs. 0.8447) were slightly lower.

**Table 2.** Ablation results. The table shows the metrics for different model variations that replace Hallmark-GCN with other frameworks such as self-attention, GAT, CNN, replace highway gate module with 3-layer MLP or use two fully connected 3-layer MLP. The best metric is bold, and the second-best metric is underlined. ^a^The model only uses the local output of the level that immediately precede it. ^b^The model uses the local output of all levels that precede it.

| Model | Accuracy | Macro-Precision | Macro-Recall | Macro-F1 |
|---|---|---|---|---|
| Hallmark-GCN + Highway<sup>a</sup> | 0.7921±0.0020 | **0.8677±0.0020** | **0.8396±0.0017** | **0.8447±0.0019** |
| Hallmark-GCN + Highway<sup>b</sup> | **0.7981±0.0042** | 0.8585±0.0039 | 0.8284±0.0042 | 0.8342±0.0043 |
| Self-attention + Highway | 0.7829±0.0058 | 0.8480±0.0028 | 0.8199±0.0033 | 0.8247±0.0029 |
| GAT + Highway | 0.7892±0.0020 | 0.8492±0.0037 | 0.8218±0.0028 | 0.8268±0.0023 |
| CNN + Highway | 0.7136±0.0101 | 0.8179±0.0103 | 0.7867±0.0095 | 0.7915±0.0102 |
| Hallmark-GCN + 3-layer MLP | 0.7636±0.0040 | 0.8614±0.0046 | 0.8164±0.0030 | 0.8263±0.0038 |
| 3-layer MLP + 3-layer MLP | 0.7368±0.0084 | 0.8363±0.0058 | 0.7937±0.0054 | 0.8029±0.0061 |

We then systematically replaced individual components of the HHRM recursive module. The Hallmark-GCN used for feature embedding was replaced with self-attention, a graph attention network (GAT), or a convolutional neural network (CNN), and the highway gate used for information filtering was replaced with a three-layer MLP. We also tested a variant in which both components were replaced by three-layer MLPs. These alternatives extract information differently. The Hallmark-GCN aggregates information using prior knowledge of gene interactions encoded in the cancer hallmark networks. Self-attention adjusts network weights through an attention mechanism. GAT adjusts network weights through attention filtered by the cancer hallmark networks. The CNN extracts information using three convolutional layers, and the MLP is a fully connected neural network for information extraction.

Compared with using only the immediately preceding level’s output (Table 2), all replacement variants showed reduced performance on all four metrics. Accuracy decreased by 0.0029-0.078, macro-precision by 0.0063-0.0498, macro-recall by 0.0178-0.0529, and macro-F1 by 0.0179-0.0532. Among the replacement variants, GAT achieved the highest accuracy (0.7892), macro-recall (0.8218), and macro-F1 (0.8268), whereas Hallmark-GCN + three-layer MLP achieved the highest macro-precision (0.8614). The CNN variant had the lowest values on all four metrics. Overall, replacing either the Hallmark-GCN or the highway gate reduced performance, most markedly for the CNN and MLP-based variants, supporting a contribution of both components within the HHRM recursive module.

### 3.3 HHRM maintains robust performance on an independent dataset

We further evaluated HHRM’s performance on a validation set of 887 samples. These samples represent a particularly challenging subset, as each was assigned a label entirely discordant with its diagnosis in the original study, yielding an accuracy of 0.0 for this subset. HHRM’s accuracy was 0.3450 and 0.3788 when evaluated against the full eight-level labels of the samples and their original levels, respectively. Model performance varied substantially across cancer types, ranging from 0.0 to 1.0 accuracy (Supplementary Excel Sheet S5). Although this performance is not satisfactory, it represents a 0.0105-0.1388 improvement on 0.24 from our previous model, which trained eight classifiers separately for each label level and then merged them to form hierarchical labels [31].

To assess generalizability, we evaluated HHRM on an external dataset called METABRIC. that contains 1,608 valid breast cancer samples with hierarchical labels of two levels, including Basal, Lum A, Lum B, and HER2, that match the subtypes in the Treehouse data (Method section 2.5). We split the data into 60% training, 20% test, and 20% validation. First, we ran HHRM and the benchmark models five times and evaluated their performance on the 20% test data. Overall, the models performed similarly on the METABRIC data, with differences ranging from 0.032 to 0.0404 across the four metrics (Table 3). No single model outperformed the others in all metrics. More specifically, accuracy ranged from 0.8907 (HARNN) to 0.9239 (HMCN_F), with a difference of 0.033. HHRM achieved an accuracy of 0.9105, ranking third between C-HMCNN (0.9221) and the 3-layer MLP (0.9047). Across the other metrics, the values ranged from 0.8877 to 0.9287 for macro-precision, from 0.8675 to 0.9079 for macro-recall, and from 0.8791 to 0.9130 for macro-F1. HHRM’s macro-precision (0.9197) was close to HMCN_F’s (0.9199), but its macro-recall (0.8675) and macro-F1 (0.8791) were the lowest among the six models. HMCN_F had the highest accuracy, macro-recall, and macro-F1, while the 3-layer MLP had the highest macro-precision.

**Table 3.** Model performance on the METABRIC dataset. The table shows the metrics for classifying four breast cancer subtypes with two levels. The best metric is bold, and the second-best metric is underlined. The metric is the average and standard deviation of the values from five runs on the 20% test data (the first four rows). The best-performing HHRM result of the five runs on the test data can be found in Supplementary Excel Sheet S3. The HHRM with the best performance is tested further on the 20% validation data and evaluated using the four metrics (the last row).

|  | Model | Accuracy | Macro-Precision | Macro-Recall | Macro-F1 |
|---|---|---|---|---|---|
| 20% test | HHRM (ours) | 0.9105±0.0081 | 0.9197±0.0166 | 0.8675±0.0183 | 0.8791±0.0221 |
|  | HARNN | 0.8907±0.0120 | 0.8877±0.0179 | 0.8917±0.0179 | 0.8881±0.0063 |
|  | HMCN_F | **0.9239±0.0062** | 0.9199±0.0201 | **0.9079±0.0145** | **0.9130±0.0061** |
|  | HMCN_R | 0.8957±0.0055 | 0.8983±0.0136 | 0.9009±0.0177 | 0.8990±0.0095 |
|  | C-HMCNN | 0.9221±0.0045 | 0.9257±0.0147 | 0.8926±0.0046 | 0.9077±0.0057 |
|  | 3-layer MLP | 0.9047±0.0045 | **0.9287±0.0132** | 0.8737±0.0077 | 0.8991±0.0098 |
| 20% val. | HHRM (ours) | 0.8659 | 0.8433 | 0.8924 | 0.8613 |

Second, we tested the best performing HHRM on the remaining 20% validation data. The model achieved comparable performance to the 20% test data, with an accuracy value of 0.8659 (Table 3). Accuracy values ranged from 0.6667 to 0.9714 across the four breast cancer subtypes (Supplementary Excel Sheet S3). Performance was lower for labels with fewer samples (Basal and HER2) than for labels with more samples (Lum A and B).

Finally, we performed zero-shot prediction on the METABRIC data, i.e., we applied the HHRM model trained on the Treehouse data directly to predict breast cancer subtypes, without retraining. The model achieved an accuracy of 0.3495 for the two-level hierarchical subtypes. This lower accuracy may reflect differences between the two datasets: METABRIC expression was measured by microarray, whereas the training data were generated by RNA sequencing, and the METABRIC labels did not exactly match those of the training data and were transformed to match its hierarchical labels.

Taken together, the model demonstrates good generalizability when using validation and independent datasets, despite having some limitations when it comes to predicting the hierarchical labels of the most challenging samples and making zero-shot predictions.

### 3.4 Effect of hierarchical complexity on model accuracy

Having shown that HHRM performs reliably in classifying hierarchical tumor subtypes, we next examined how the complexity of the subtype hierarchy relates to its performance. The pan-cancer data comprises complex hierarchical tumor subtypes, which pose a classification challenge characterized by intricate intra-tumor and inter-tumor heterogeneity. We therefore analyzed how tumor subtypes with different numbers of label combinations and hierarchical levels relate to the model’s performance. To this end, we defined a difficulty coefficient, *comb* / (*max_level* × *sample_size*), to denote the prediction difficulty of each cancer type considering all labels at eight levels. Here, *comb* is the number of label combinations in a cancer type, computed as the product of the number of labels at each level, *max_level* is the total number of hierarchical levels of the cancer type and has a value of 8, and *sample_size* is the number of training samples of the cancer type. The coefficient increases with the cancer type’s number of labels across all levels and decreases as the number of samples available for model training increases.

The coefficient showed a significant negative correlation with model accuracy on the test data (Spearman correlation coefficient = −0.437 with a p-value of 0.0256; Figure 4). Model performance declined as the coefficient increased, but the extent of the decline depended on each cancer type’s label combinations and sample size. Specifically, the model achieved 100% accuracy for Ewing sarcoma (T005 EWING) and testicular germ cell tumor, seminoma (T025 TGCT SEM), both of which have no subtypes and are represented by one label at one level. In contrast, accuracy dropped to 0.5556 for ovarian cystadenocarcinoma (T023 OV), which has 56,953,125 label combinations across three levels and only 451 samples for model training. Cancer types with a coefficient at a similar level to OV but with more labels showed higher accuracy: leukemia (T005 LEUK), with 302,603,400 label combinations and 888 samples, and squamous cell or bladder carcinoma (T012 SCC/BLCA), with 729,797,600 label combinations and 1,657 samples, reached accuracies of 0.9181 and 0.7964, respectively, with substantially more samples than OV.

**Figure 4.**
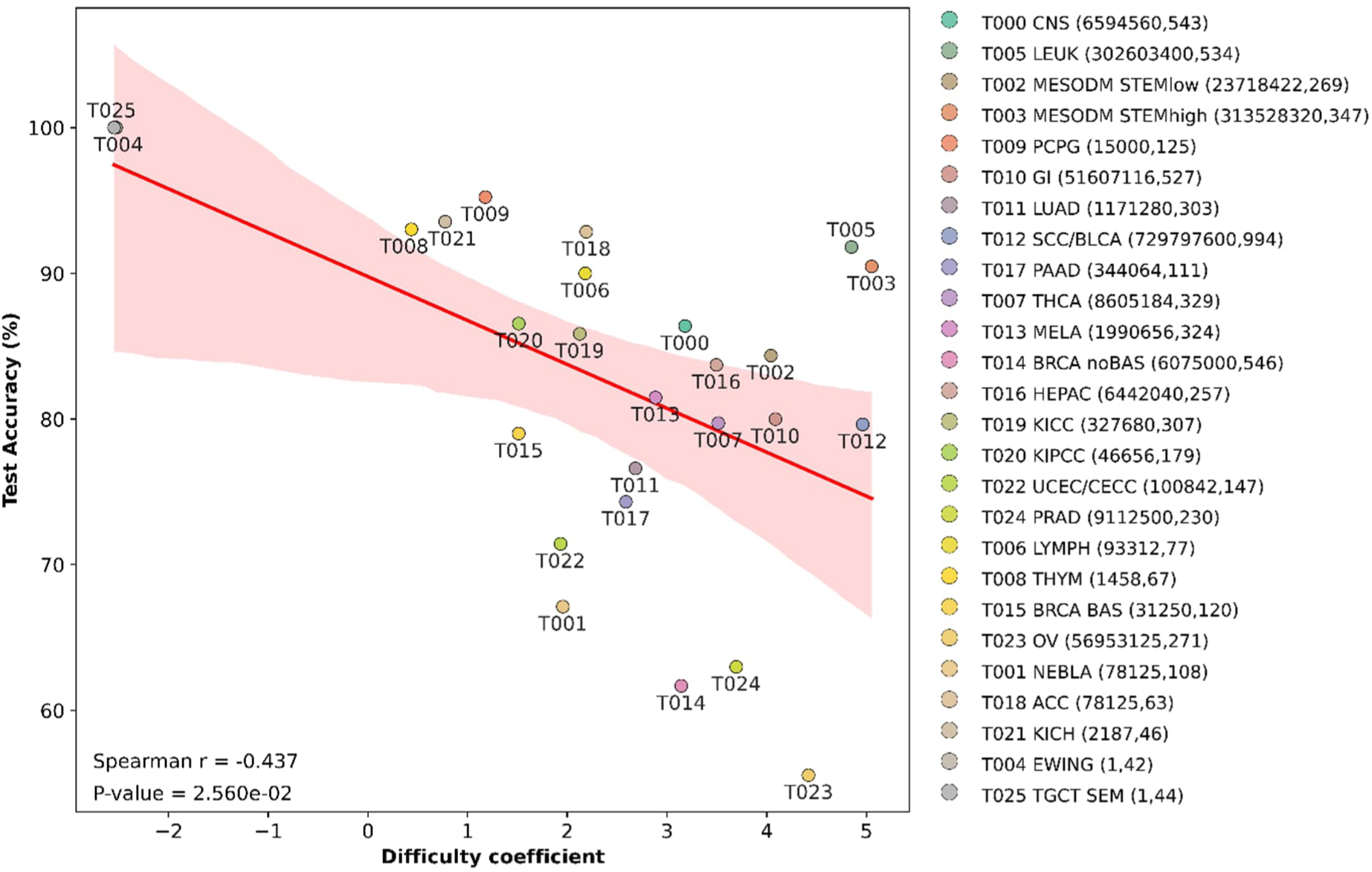
Association between label complexity and model performance across the 26 cancer types. Each circle represents a cancer type with the corresponding model accuracy on y-axis and difficult coefficient on x-axis (log10 transformed). The line is a linear regression of all circles, and the shaded area indicates the 95% confidence interval for the fitting line. The numbers in the paratheses indicate the number of possible label combinations and the total number of training samples for each cancer type, respectively. The data for drawing plot can be found in Supplementary Excel Sheet S5.

Taken together, higher heterogeneity in tumor subtypes, as reflected by the number of labels at each level, was associated with lower classifier performance. The number of samples available for model training may compensate for this decline, although its impact depended on the type of cancer.

### 3.5 Functional characterization of genes prioritized by SHAP analysis

The recursive modules of HHRM use a GCN to integrate gene expression profiles with cancer hallmark networks for feature embedding, and the embedded features are used to classify hierarchical tumor subtypes. This design allowed us to combine machine learning with consensus biological knowledge to interpret model decisions at both hallmark and gene levels. We first computed SHAP values for hallmark genes at each hierarchical level and ranked the genes at each level by their average absolute SHAP values; genes with higher values were regarded as more important for the model (METHOD Section 2.7). We then extracted the top 10 genes at each level, which resulted in 38 unique genes across the ten cancer hallmarks (Figure 5 top). Of these genes, 17 were associated with only one cancer hallmark, and 21 were shared by multiple hallmarks. Most genes were associated with hallmarks related to maintenance of cell proliferation, evasion of growth suppressors, and activation of invasion and metastasis.

**Figure 5.**
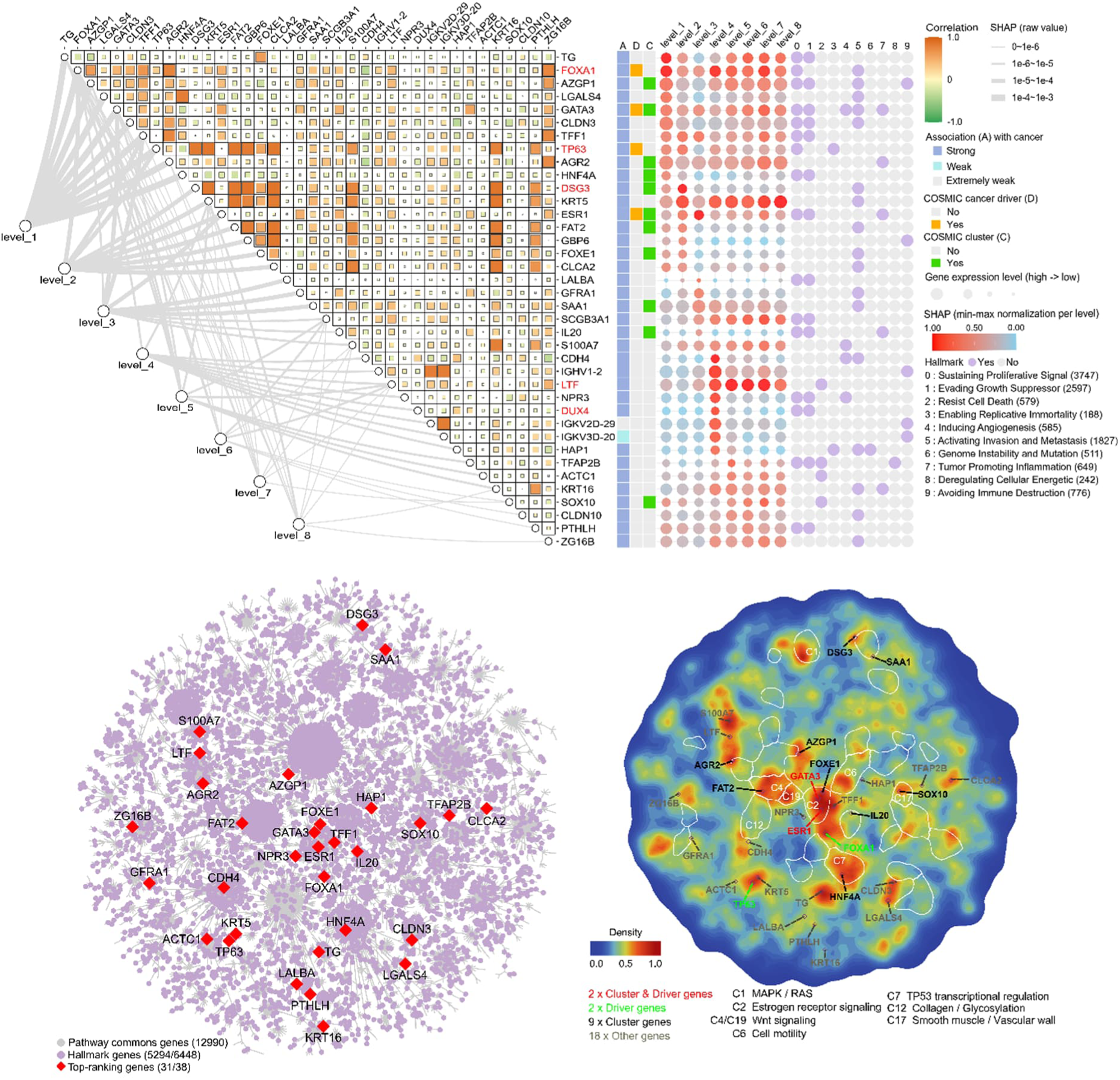
Model interpretation using top genes ranked by SHAP values. Top: The heatmap displays Pearson correlation coefficients among the top 10 ranking genes at each hierarchical level. The gray lines connecting hierarchical levels and genes show the genes’ membership within each label level. Line width represents each gene’s SHAP value. On the right annotations, the boxes indicate the category of genes’ associations (A) with cancer by searching for evidence from the literature (Supplementary Excel Sheet S6). Dark blue boxes indicate genes for which there is substantial evidence of a strong association with specific cancers (e.g., genes that drive or suppress cancer progression). Light blue boxes indicate genes that may be associated with certain cancers, but the evidence is incomplete and has not reached a research consensus. Gray boxes indicate genes that are not documented as being related to any cancer, but that may be expressed in normal tissue. The subsequent squares whether indicate whether a gene belongs to COSMIC cluster (C) or COSMIC cancer driver genes (D). The dots’ color code and sizes represent each gene’s normalized SHAP value at each subtype level and the average gene expression level in all samples, respectively. The SHAP values of the genes are normalized using the min-max method. Higher values indicate greater importance at a specific level. The genes highlighted in red are the ones whose SHAP profiles are described in the texts. The further dots indicate genes’ membership among the ten cancer hallmarks (purple denotes a membership otherwise gray). **Bottom left**: The background gene regulatory network comprises 12,990 genes from Pathway Commons (grey). The highlighted genes include hallmark genes (purple) and the top-ranking genes (blue) in the heat map. **Bottom right**: The signal aggregation density map illustrates the network-based aggregation of gene signals. SHAP values were assigned as signals to the hallmark genes (highlighted in pink and red in the bottom left panel), while all other genes in the background network were set to zero. Each signal was then propagated across the network according to its topology, with signal decay following the default Weibull decay function. For each point in the two-dimensional image space, the propagated signals from neighboring genes were aggregated into a scalar using mean aggregation, which defines the density value at that point. Darker red indicates higher aggregated signal density, while darker blue indicates lower signal density. The white circles correspond to the 30 COSMIC clusters (Supplementary Figure S4 and Supplementary Excel Sheet S7). The highlighted clusters are eight COSMIC gene clusters that visibly overlap with hallmark gene clusters, including C1, C2, C4, C6, C7, C12, C17, and C19, indicating that hallmark genes show high aggregated signal density in these clusters. The highlighted genes are the top-ranking genes. Gene names are color-coded according to their classification: red for both COSMIC cluster and driver genes; black for COSMIC cluster genes only; green for COSMIC cancer driver genes only; and grey for genes not belonging to any set.

To directly compare the hallmarks, we aggregated the SHAP values of the genes in each hallmark to create a hallmark SHAP profile at each hierarchical level (Figure S5). Across all levels, the hallmarks related to angiogenesis and tissue invasion and metastasis contributed the most to the classification of hierarchical tumor subtypes, while the hallmarks related to genome instability and replicative immortality contributed the least. Thus, the model relied most heavily on hallmarks describing the tumor microenvironment and invasive behavior and relatively little on hallmarks describing intrinsic cellular maintenance.

To provide context for the SHAP-ranked genes in relation to known cancer genes, we used information from the COSMIC database, which contains 554 tier 1 cancer driver genes [40]. These genes are well-established cancer genes for which there is robust experimental and published evidence that mutations alter the activity of the gene product to drive oncogenic transformation [42]. We identified four COSMIC cancer driver genes (FOXA1, GATA3, ESR1, and TP63) from the top-ranking genes and they have higher SHAP values at levels 1-3 (mean = 0.61 vs. 0.30; two-sided Mann-Whitney U test p-value = 0.001) and higher overall SHAP values (mean = 0.60 vs. 0.36; p-value = 0.017) than other genes (Figure 5 top). More specifically, FOXA1 is highly expressed and shows importance at all levels (SHAP ≥ 0.5) and peaks at levels 1 (SHAP = 0.99), 4 (SHAP = 0.98), and 7 (SHAP = 0.97). FOXA1 is involved in sustaining proliferative signaling by enabling estrogen receptor activity [43]. High SHAP values at all levels are consistent with FOXA1’s role in defining luminal-type epithelial lineages in breast and prostate cancers [44], [45], while also suggesting that FOXA1 can contribute to finer distinctions between tumor subtypes. TP63 is moderately expressed and is important at the primary and secondary levels (SHAP = 0.62 and 0.72, respectively) but declines at higher levels. The gene plays a role in sustaining proliferative signaling, evading growth suppressors, and resisting cell death [46]. The higher SHAP value at the primary and secondary levels indicates a more significant role in squamous differentiation than in refined distinction in tumor subtypes.

We next examined genes that are not COSMIC cancer driver genes. DSG3, which is moderately expressed, has its highest SHAP value at level 2 (SHAP = 1.0) and much lower values at other levels, suggesting that it mainly helps separate coarse tumor types rather than refined ones. The moderately expressed DUX4 has a high SHAP value at level 4 (0.93), alongside the immunoglobulin variable-region genes IGHV1-2 (SHAP = 0.97), IGKV2D-29 (SHAP = 0.87), and IGKV3D-20 (SHAP = 0.86). This suggests DUX4’s important role in classifying tumor subtypes at one specific level. LTF is highly expressed and has high SHAP values at levels 5-8, suggesting that it contributes mainly to classifying refined tumor subtypes rather than coarse tumor types.

Of note, several top-ranking genes, including TP63, DSG3, KRT5, FAT2, GBP6, CLCA2, S100A7, and IGHV1-2, exhibited strong correlation in expression with other top-ranking genes. As indicated by the SHAP profiles, their importance differs across levels. TP63 ranks among the top genes at levels 1 and 2 and is strongly correlated with five genes, whereas DSG3 only ranks top at level 2 and is strongly correlated with eight genes. Since correlated genes can share predictive information, these genes may be partly interchangeable for the classifier. This suggests redundancy in transcriptomic information of genes for hierarchical classification.

Furthermore, we used a network-based tool, PathwaySpace [18], to analyze the overlap between the SHAP-ranked genes and COSMIC cancer driver genes, as well as their involvement in cancer driver genes’ clusters (METHOD Section 2.7). The tool mapped the hallmark genes onto the gene regulatory network from Pathway Commons. 5,294 of the 6,448 hallmark genes (82.1%) and 31 of the 38 SHAP-ranked genes (81.6%) were found in the network (Figure 5 bottom left). The hallmark genes formed clusters that overlapped with and differed from the 30 COSMIC gene clusters (Figure 5 bottom right). They mainly overlapped with eight COSMIC gene clusters, namely those associated with the MAPK/RAS pathway (C1), estrogen receptor signaling (C2), WNT signaling (C4 and C19), cell motility (C6), TP53 transcriptional regulation (C7), collagen C glycosylation (C12), and smooth muscle vascular wall (C17).

Among the 38 SHAP-ranked genes, 31 were found in the gene regulatory network (Figure 5 bottom left). Compared to the other hallmark genes, the top five ranked genes in each level were enriched in COSMIC cancer driver genes, with an odds ratio of 4.87 (Fisher’s exact test p-value = 0.0156) (Supplementary Table S2). Of the 31 genes, two (ESR1 and GATA3) were cancer driver genes that also fell into a cluster, two others (FOXA1 and TP63) were cancer driver genes only, and nine fell into COSMIC gene clusters. ESR1 and GATA3 are strongly associated with tumorigenesis and progression in breast cancer [48]. Together with FOXA1, they form a transcriptional axis that defines luminal mammary epithelial identity in breast cancer [49]. This axis operates through a hierarchical chromatin-remodeling mechanism in which GATA3 and FOXA1 prepare enhancer accessibility for ligand-activated ESR1. The co-option or mutation of these proteins in tumors reshapes the transcriptional programs that govern growth, endocrine response, and metastasis [50], [51]. The driver gene TP63 encodes multiple isoforms and rarely undergoes mutation in cancers [52]; instead, it is amplified or overexpressed, which confers a proliferative advantage and sustains squamous identity across multiple tumor types [47]. The nine genes within the COSMIC gene clusters could be grouped into four categories: lineage transcriptional control (FOXE1, HNF4A, SOX10) [53], [54], [55], cell-adhesion and surface signaling (DSG3, FAT2) [56], [57], secreted or extracellular microenvironmental regulators (AZGP1, IL20, SAA1) [58], [59], [60], and epithelial homeostasis and inflammation (AGR2) [61]. Although the remaining 18 genes do not belong to the COSMIC driver genes and clusters, they have also been reported to be related to cancer through literature search (Supplementary Excel Sheet S6).

Taken together, the identified top-ranking genes include those that overlap with known tumor driver genes that have been functionally annotated, as well as under-investigated genes that may require further study in the context of cancer.

## Conclusions and Discussion

We present the first model that can classify hierarchical tumor subtypes considering the inherent sequential dependence in the tumor subtypes. The model achieves an accuracy of nearly 0.80 for hierarchical label prediction, which is notable given the considerable complexity of the classification task. This task spans 1,713 tumor subtypes across eight hierarchical levels with pronounced label imbalance. The model’s architecture is inspired by a recurrent neural network for classifying hierarchical documents [33]. The resulting model can precisely classify a sample’s subtype level by level via integrating the sample’s transcriptomic profile and the hierarchical structure in tumor subtypes. Similar ideas can be found for improving protein function prediction [26] and metagenome-assembled genomes [29]. This demonstrates that the organized relationship among labels is an informative feature for hierarchical multi-label classification.

Interestingly, we expect that incorporating all preceding local outputs, rather than only the immediately preceding one, would provide the current level’s module with richer information, thereby improving performance since more information would be passed forward from earlier to later levels. However, the variant that used only the immediately preceding level’s output achieved comparable performance. This suggests that the immediately preceding level’s output encodes sufficient hierarchical information to predict the current level. This is likely because each level’s prediction is derived recursively from all earlier levels. This means that the prediction of the level immediately preceding it is the primary conditioning variable, and earlier labels are often indirectly represented through it [30]. Therefore, explicitly reintroducing the full history of local outputs adds limited additional information.

We use prior biological knowledge to filter the genome-wide transcriptome and use cancer hallmark genes as the model’s input features. Unlike purely data-driven feature selection, this approach restricts the model to genes with established biological relevance to cancer, making the model’s predictions more interpretable [62]. This is demonstrated with the model’s performance on an independent breast cancer cohort that has four subtypes with two levels. After retraining the model with 60% data, it reaches the accuracy greater than 91% for test and validation data. However, when performing on the same data without retraining, the model’s accuracy drops to 0.3495. This limitation is due to the lack of an independent dataset with a comparable hierarchical tumor structure for external validation. This restricts our capacity to adequately evaluate the model’s generalizability beyond the existing cohort. One potential solution would be to generate a synthetic independent validation cohort, but it still faces the challenge to preserve clinical validity [63].

Finally, by combining SHAP with a network-based enrichment analysis, we elaborate the role of the identified important gene in cancer. We found that the gene list not only overlaps with known cancer driver genes significantly but also contains understudied genes worth investigating in the future. However, such a post hoc approach does not integrate prior biological knowledge into the model’s architecture limiting its interpretability to trace back to specific molecular mechanisms and biological processes in cancer subtypes. Closing this gap requires a model that treats hierarchy and interpretability as primary design constraints [64], [65]. Although models that embed prior biological knowledge in their architecture may offer more interpretable explanations, their performance gains do not necessarily reflect the value of that knowledge. These gains may instead come from the sparsity-induced regularization imposed by the knowledge-derived structure [66].

In conclusion, HHRM’s ability to stratify intricate hierarchical tumor subtypes while maintaining interpretability represents a significant methodological advancement, holding promise for precision diagnostics in cancer.

## AI usage declaration

After drafting the manuscript, we used Claude Sonnet 5.5 to check the language and correct the grammar. We take full responsibility for the text refined by the AI-writing tool.

## Ethics, Consent to Participate, and Consent to Publish declarations

not applicable.

## Data availability

The code used in this study is accessible at https://github.com/laixn/HHRM for non-commercial use. The data and model weights are available at https://doi.org/10.5281/zenodo.23589666.

## Author contributions

Ǫingsong Zhang (Data curation [lead], Formal analysis [supporting], Methodology [equal], Software [lead], Visualization [equal], Writing—original draft [supporting], Writing—review C editing [supporting]), Fei Liu (Funding acquisition [equal], Methodology [supporting], Project administration [supporting], Resources [equal], Supervision [supporting], Writing—review C editing [supporting]), and Xin Lai (Conceptualization [lead], Formal analysis [lead], Funding acquisition [equal], Investigation [lead], Methodology [Equal], Project administration [lead], Resources [equal], Software [supporting], Supervision [lead], Validation [lead], Visualization [equal], Writing—original draft [lead], Writing—review C editing [lead])

## References

[1] Cancer Genome Atlas Research Network et al., “The Cancer Genome Atlas Pan-Cancer analysis project,” Nat Genet, vol. 45, no. 10, pp. 1113–1120, Oct. 2013, doi: 10.1038/ng.2764.

[2] K. A. Hoadley et al., “Cell-of-Origin Patterns Dominate the Molecular Classification of 10,000 Tumors from 33 Types of Cancer,” Cell, vol. 173, no. 2, pp. 291–304.e6, Apr. 2018, doi: 10.1016/j.cell.2018.03.022.

[3] J. Guinney et al., “The consensus molecular subtypes of colorectal cancer,” Nat Med, vol. 21, no. 11, pp. 1350–1356, Nov. 2015, doi: 10.1038/nm.3967.

[4] L. Zhao, V. H. F. Lee, M. K. Ng, H. Yan, and M. F. Bijlsma, “Molecular subtyping of cancer: current status and moving toward clinical applications,” Brief Bioinform, vol. 20, no. 2, pp. 572–584, Mar. 2019, doi: 10.1093/bib/bby026.

[5] F. Bertucci, P. Finetti, A. Goncalves, and D. Birnbaum, “The therapeutic response of ER+/HER2-breast cancers differs according to the molecular Basal or Luminal subtype,” NPJ Breast Cancer, vol. 6, p. 8, 2020, doi: 10.1038/s41523-020-0151-5.

[6] J. D. Hainsworth and F. A. Greco, “Cancer of Unknown Primary Site: New Treatment Paradigms in the Era of Precision Medicine,” Am Soc Clin Oncol Educ Book, vol. 38, pp. 20–25, May 2018, doi: 10.1200/EDBK_100014.

[7] N. Pavlidis and G. Pentheroudakis, “Cancer of unknown primary site,” Lancet, vol. 379, no. 9824, pp. 1428–1435, Apr. 2012, doi: 10.1016/S0140-6736(11)61178-1.

[8] D. Feldner-Busztin et al., “Dealing with dimensionality: the application of machine learning to multi-omics data,” Bioinformatics, vol. 39, no. 2, p. btad021, Feb. 2023, doi: 10.1093/bioinformatics/btad021.

[9] C.-Y. Hsu et al., “AI-driven multi-omics integration in precision oncology: bridging the data deluge to clinical decisions,” Clin Exp Med, vol. 26, no. 1, p. 29, Nov. 2025, doi: 10.1007/s10238-025-01965-9.

[10] F. Siddique, M. Shehata, M. Ghazal, S. Contractor, and A. El-Baz, “Lung Cancer Subtyping: A Short Review,” Cancers (Basel), vol. 16, no. 15, p. 2643, Jul. 2024, doi: 10.3390/cancers16152643.

[11] Y. Li et al., “A comprehensive genomic pan-cancer classification using The Cancer Genome Atlas gene expression data,” BMC Genomics, vol. 18, no. 1, p. 508, Jul. 2017, doi: 10.1186/s12864-017-3906-0.

[12] W. Jiao et al., “A deep learning system accurately classifies primary and metastatic cancers using passenger mutation patterns,” Nat Commun, vol. 11, no. 1, p. 728, Feb. 2020, doi: 10.1038/s41467-019-13825-8.

[13] D. Yuan et al., “crossNN is an explainable framework for cross-platform DNA methylation-based classification of tumors,” Nat Cancer, vol. 6, no. 7, pp. 1283– 1294, Jul. 2025, doi: 10.1038/s43018-025-00976-5.

[14] F. Chen, D. S. Chandrashekar, S. Varambally, and C. J. Creighton, “Pan-cancer molecular subtypes revealed by mass-spectrometry-based proteomic characterization of more than 500 human cancers,” Nat Commun, vol. 10, no. 1, p. 5679, Dec. 2019, doi: 10.1038/s41467-019-13528-0.

[15] S. Cascianelli et al., “Multi-label transcriptional classification of colorectal cancer reflects tumor cell population heterogeneity,” Genome Med, vol. 15, no. 1, p. 37, May 2023, doi: 10.1186/s13073-023-01176-5.

[16] J. Vibert et al., “Identification of Tissue of Origin and Guided Therapeutic Applications in Cancers of Unknown Primary Using Deep Learning and RNA Sequencing (TransCUPtomics),” J Mol Diagn, vol. 23, no. 10, pp. 1380–1392, Oct. 2021, doi: 10.1016/j.jmoldx.2021.07.009.

[17] J. Michuda et al., “Validation of a Transcriptome-Based Assay for Classifying Cancers of Unknown Primary Origin,” Mol Diagn Ther, vol. 27, no. 4, pp. 499–511, Jul. 2023, doi: 10.1007/s40291-023-00650-5.

[18] K. Ellrott et al., “Classification of non-TCGA cancer samples to TCGA molecular subtypes using compact feature sets,” Cancer Cell, vol. 43, no. 2, pp. 195–212.e11, Feb. 2025, doi: 10.1016/j.ccell.2024.12.002.

[19] F. Comitani et al., “Diagnostic classification of childhood cancer using multiscale transcriptomics,” Nat Med, vol. 29, no. 3, pp. 656–666, Mar. 2023, doi: 10.1038/s41591-023-02221-x.

[20] Y. Yang et al., “Hierarchical classification-based pan-cancer methylation analysis to classify primary cancer,” BMC Bioinformatics, vol. 24, no. 1, p. 465, Dec. 2023, doi: 10.1186/s12859-023-05529-0.

[21] C. de Margerie-Mellon and G. Chassagnon, “Artificial intelligence: A critical review of applications for lung nodule and lung cancer,” Diagn Interv Imaging, vol. 104, no. 1, pp. 11–17, Jan. 2023, doi: 10.1016/j.diii.2022.11.007.

[22] P. Pati et al., “Hierarchical graph representations in digital pathology,” Med Image Anal, vol. 75, p. 102264, Jan. 2022, doi: 10.1016/j.media.2021.102264.

[23] A. M. Rinaldi, C. Russo, and C. Tommasino, “Hierarchical graph networks for breast cancer subtype classification,” Data Min Knowl Disc, vol. 40, no. 6, p. 104, Sep. 2026, doi: 10.1007/s10618-026-01259-y.

[24] B. Li and S. Nabavi, “A multimodal graph neural network framework for cancer molecular subtype classification,” BMC Bioinformatics, vol. 25, no. 1, p. 27, Jan. 2024, doi: 10.1186/s12859-023-05622-4.

[25] J. Wehrmann, R. Cerri, and R. Barros, “Hierarchical Multi-Label Classification Networks,” in Proceedings of the 35th International Conference on Machine Learning, PMLR, Jul. 2018, pp. 5075–5084.

[26] R. Cerri, R. C. Barros, A. C. P L F de Carvalho, and Y. Jin, “Reduction strategies for hierarchical multi-label classification in protein function prediction,” BMC Bioinformatics, vol. 17, no. 1, p. 373, Sep. 2016, doi: 10.1186/s12859-016-1232-1.

[27] E. Giunchiglia and T. Lukasiewicz, “Coherent Hierarchical Multi-Label Classification Networks,” in Advances in Neural Information Processing Systems, Curran Associates, Inc., 2020, pp. 9662–9673.

[28] J. Mikkonen, T. J. Rintala, and V. Fortino, “Hierarchical modeling of tumor subtypes in cell lines using large-scale genomic datasets,” iScience, vol. 29, no. 9, p. 117315, Sep. 2026, doi: 10.1016/j.isci.2026.117315.

[29] S. Kutuzova et al., “Improving metagenome binning by integrating intrinsic features and taxonomy,” Nat Biotechnol, Apr. 2026, doi: 10.1038/s41587-026-03098-0.

[30] C. N. Silla and A. A. Freitas, “A survey of hierarchical classification across different application domains,” Data Min Knowl Disc, vol. 22, no. 1, pp. 31–72, Jan. 2011, doi: 10.1007/s10618-010-0175-9.

[31] Ǫ. Zhang, F. Liu, and X. Lai, “HallmarkGraph: a cancer hallmark informed graph neural network for classifying hierarchical tumor subtypes,” Bioinformatics, vol. 41, no. 9, p. btaf444, Sep. 2025, doi: 10.1093/bioinformatics/btaf444.

[32] M. Milacic et al., “The Reactome Pathway Knowledgebase 2024,” Nucleic Acids Res, vol. 52, no. D1, pp. D672–D678, Jan. 2024, doi: 10.1093/nar/gkad1025.

[33] W. Huang et al., “Hierarchical Multi-label Text Classification: An Attention-based Recurrent Network Approach,” in Proceedings of the 28th ACM International Conference on Information and Knowledge Management, in CIKM ‘19. New York, NY, USA: Association for Computing Machinery, Nov. 2019, pp. 1051–1060. doi: 10.1145/3357384.3357885.

[34] R. K. Srivastava, K. Greff, and J. Schmidhuber, “Training Very Deep Networks,” in Advances in Neural Information Processing Systems, Curran Associates, Inc., 2015.

[35] B. Pereira et al., “The somatic mutation profiles of 2,433 breast cancers refines their genomic and transcriptomic landscapes,” Nat Commun, vol. 7, p. 11479, May 2016, doi: 10.1038/ncomms11479.

[36] O. M. Rueda et al., “Dynamics of breast-cancer relapse reveal late-recurring ER-positive genomic subgroups,” Nature, vol. 567, no. 7748, pp. 399–404, Mar. 2019, doi: 10.1038/s41586-019-1007-8.

[37] C. Curtis et al., “The genomic and transcriptomic architecture of 2,000 breast tumours reveals novel subgroups,” Nature, vol. 486, no. 7403, pp. 346–352, Apr. 2012, doi: 10.1038/nature10983.

[38] S. M. Lundberg and S.-I. Lee, “A Unified Approach to Interpreting Model Predictions,” in Advances in Neural Information Processing Systems, Curran Associates, Inc., 2017.

[39] I. Rodchenkov et al., “Pathway Commons 2019 Update: integration, analysis and exploration of pathway data,” Nucleic Acids Res, vol. 48, no. D1, pp. D489–D497, Jan. 2020, doi: 10.1093/nar/gkz946.

[40] Z. Sondka et al., “COSMIC: a curated database of somatic variants and clinical data for cancer,” Nucleic Acids Res, vol. 52, no. D1, pp. D1210–D1217, Jan. 2024, doi: 10.1093/nar/gkad986.

[41] G. Huang, Z. Liu, L. Van Der Maaten, and K. Ǫ. Weinberger, “Densely Connected Convolutional Networks,” in 2017 IEEE Conference on Computer Vision and Pattern Recognition (CVPR), Jul. 2017, pp. 2261–2269. doi: 10.1109/CVPR.2017.243.

[42] Z. Sondka, S. Bamford, C. G. Cole, S. A. Ward, I. Dunham, and S. A. Forbes, “The COSMIC Cancer Gene Census: describing genetic dysfunction across all human cancers,” Nat Rev Cancer, vol. 18, no. 11, pp. 696–705, Nov. 2018, doi: 10.1038/s41568-018-0060-1.

[43] A. Hurtado, K. A. Holmes, C. S. Ross-Innes, D. Schmidt, and J. S. Carroll, “FOXA1 is a key determinant of estrogen receptor function and endocrine response,” Nat Genet, vol. 43, no. 1, pp. 27–33, Jan. 2011, doi: 10.1038/ng.730.

[44] E. J. Adams et al., “FOXA1 mutations alter pioneering activity, differentiation and prostate cancer phenotypes,” Nature, vol. 571, no. 7765, pp. 408–412, Jul. 2019, doi: 10.1038/s41586-019-1318-9.

[45] J. Jeong, et al., “A FOXA1/SPDEF Co-regulatory Axis Drives ERBB2 and Suppresses TEAD/YAP-Driven EMT to Maintain Luminal Identity in HER2+ Breast Cancer,” Oct. 09, 2025, bioRxiv. doi: 10.1101/2024.04.16.589460.

[46] M. A. Moses et al., “Molecular Mechanisms of p63-Mediated Squamous Cancer Pathogenesis,” Int J Mol Sci, vol. 20, no. 14, p. 3590, Jul. 2019, doi: 10.3390/ijms20143590.

[47] R. Pecorari, F. Bernassola, G. Melino, and E. Candi, “Distinct interactors define the p63 transcriptional signature in epithelial development or cancer,” Biochem J, vol. 479, no. 12, pp. 1375–1392, Jun. 2022, doi: 10.1042/BCJ20210737.

[48] L. Porras, H. Ismail, and S. Mader, “Positive Regulation of Estrogen Receptor Alpha in Breast Tumorigenesis,” Cells, vol. 10, no. 11, p. 2966, Oct. 2021, doi: 10.3390/cells10112966.

[49] E. M. Martin, K. A. Orlando, K. Yokobori, and P. A. Wade, “The estrogen receptor/GATA3/FOXA1 transcriptional network: lessons learned from breast cancer,” Curr Opin Struct Biol, vol. 71, pp. 65–70, Dec. 2021, doi: 10.1016/j.sbi.2021.05.015.

[50] V. Theodorou, R. Stark, S. Menon, and J. S. Carroll, “GATA3 acts upstream of FOXA1 in mediating ESR1 binding by shaping enhancer accessibility,” Genome Res, vol. 23, no. 1, pp. 12–22, Jan. 2013, doi: 10.1101/gr.139469.112.

[51] Y. Zekri et al., “Comprehensive functional annotation of ESR1-driven enhancers in breast cancer reveals hierarchical activity independent of genomic and epigenomic contexts,” Genome Res, vol. 35, no. 7, pp. 1530–1543, Jul. 2025, doi: 10.1101/gr.280320.124.

[52] G. Melino, X. Lu, M. Gasco, T. Crook, and R. A. Knight, “Functional regulation of p73 and p63: development and cancer,” Trends Biochem Sci, vol. 28, no. 12, pp. 663– 670, Dec. 2003, doi: 10.1016/j.tibs.2003.10.004.

[53] S. C. Credendino et al., “FOXE1-Dependent Regulation of Macrophage Chemotaxis by Thyroid Cells In Vitro and In Vivo,” Int J Mol Sci, vol. 22, no. 14, p. 7666, Jul. 2021, doi: 10.3390/ijms22147666.

[54] D.-D. Lv, L.-Y. Zhou, and H. Tang, “Hepatocyte nuclear factor 4α and cancer-related cell signaling pathways: a promising insight into cancer treatment,” Exp Mol Med, vol. 53, no. 1, pp. 8–18, Jan. 2021, doi: 10.1038/s12276-020-00551-1.

[55] X. Lai et al., “SOX10, MITF, and microRNAs: Decoding their interplay in regulating melanoma plasticity,” Int J Cancer, vol. 157, no. 7, pp. 1277–1293, Oct. 2025, doi: 10.1002/ijc.35499.

[56] I. O. Ilie et al., “Immunoprofile of some transmembrane adhesion molecules in oral squamous cell carcinoma,” Rom J Morphol Embryol, vol. 66, no. 3, pp. 463–478, 2025, doi: 10.47162/RJME.66.3.04.

[57] M. Govindarajan et al., “Pan-cancer N-glycoproteomic atlas of patient-derived xenografts uncovers FAT2 as an actionable surface target,” Cell Rep Med, vol. 7, no. 2, p. 102578, Feb. 2026, doi: 10.1016/j.xcrm.2025.102578.

[58] C. Burdelski et al., “Reduced AZGP1 expression is an independent predictor of early PSA recurrence and associated with ERG-fusion positive and PTEN deleted prostate cancers,” Int J Cancer, vol. 138, no. 5, pp. 1199–1206, Mar. 2016, doi: 10.1002/ijc.29860.

[59] V. Maggisano et al., “IL-20 Subfamily Biological Effects: Mechanistic Insights and Therapeutic Perspectives in Cancer,” Int J Mol Sci, vol. 26, no. 15, p. 7320, Jul. 2025, doi: 10.3390/ijms26157320.

[60] Y. Chang, Y. Liu, Y. Zou, and R. D. Ye, “Recent Advances in Studies of Serum Amyloid A: Implications in Inflammation, Immunity and Tumor Metastasis,” Int J Mol Sci, vol. 26, no. 3, p. 987, Jan. 2025, doi: 10.3390/ijms26030987.

[61] E. Boisteau et al., “Anterior gradient proteins in gastrointestinal cancers: from cell biology to pathophysiology,” Oncogene, vol. 41, no. 42, pp. 4673–4685, Oct. 2022, doi: 10.1038/s41388-022-02452-1.

[62] M. Rezapour, S. V. Trefry, L. A. Opoku, and A. Narayanan, “Artificial Intelligence in Bulk RNA-Seq: Challenges and Potential Solutions,” Comput Struct Biotechnol J, vol. 35, no. 1, p. 0039, 2026, doi: 10.34133/csbj.0039.

[63] A. Koul, D. Duran, and T. Hernandez-Boussard, “Synthetic data, synthetic trust: navigating data challenges in the digital revolution,” Lancet Digit Health, vol. 7, no. 11, p. 100924, Nov. 2025, doi: 10.1016/j.landig.2025.100924.

[64] M. Wysocka, O. Wysocki, M. Zufferey, D. Landers, and A. Freitas, “A systematic review of biologically-informed deep learning models for cancer: fundamental trends for encoding and interpreting oncology data,” BMC Bioinformatics, vol. 24, no. 1, p. 198, May 2023, doi: 10.1186/s12859-023-05262-8.

[65] J. Duan et al., “Discovering proteo-transcriptomic networks via biologically informed heterogeneous graph learning,” Nucleic Acids Res, vol. 54, no. 8, p. gkag386, Apr. 2026, doi: 10.1093/nar/gkag386.

[66] I. Caranzano et al., “Sparsity is all you need: rethinking biologically informed neural networks,” Brief Bioinform, vol. 27, no. 4, p. bbag425, Jul. 2026, doi: 10.1093/bib/bbag425.

